# Optical Tissue Clearing Enables Rapid, Precise and Comprehensive Assessment of Three-Dimensional Morphology in Experimental Nerve Regeneration Research

**DOI:** 10.1101/2021.01.28.428623

**Authors:** Simeon C. Daeschler, Jennifer Zhang, Tessa Gordon, Gregory H. Borschel

**Affiliations:** SickKids Research Institute, Neuroscience and Mental Health Program, Toronto, Ontario, Canada; Institute of Biomaterials and Biomedical Engineering, University of Toronto, Toronto, Ontario, Canada; Division of Plastic and Reconstructive Surgery, the Hospital for Sick Children, Toronto, Ontario, Canada; Indiana University School of Medicine, Indianapolis, Indiana, USA

**Keywords:** Peripheral nerve, nerve regeneration, tissue clearing, morphometry, three-dimensional imaging, image segmentation

## Abstract

Morphological analyses are key outcome assessments for nerve regeneration studies but are historically limited to tissue sections. Novel optical tissue clearing techniques enabling three-dimensional imaging of entire organs at a subcellular resolution have revolutionized morphological studies of the brain. To extend their applicability to experimental nerve repair studies we adapted these techniques to nerves and their motor and sensory targets in rats. The solvent-based protocols rendered harvested peripheral nerves and their target organs transparent within 24 h while preserving tissue architecture and fluorescence. Optical clearing is compatible with conventional laboratory techniques, including retrograde labelling studies, and computational image segmentation, providing fast and precise cell quantitation. Further, optically cleared organs enable three-dimensional morphometry at an unprecedented scale including dermatome-wide innervation studies, tracing of intramuscular nerve branches and mapping of neurovascular networks. Given their wide-ranging applicability, rapid processing times and low costs, tissue clearing techniques are likely to be a key technology for next-generation nerve repair studies.

## Introduction

Experimental nerve regeneration studies often rely on microscopic morphological analysis of nerve tissue and target organs to assess experimental outcomes. Traditionally, morphological analysis requires tissue sectioning to enable microscopic examination because light scattering limits the sampling depth to a few micrometers. Although tissue sectioning has been successfully used throughout decades of biomedical research, it disrupts the three-dimensional tissue architecture and bears the risk of cutting artifacts. Moreover, large organs such as spinal cord or skeletal muscle in adult rat models often generate hundreds of serial tissue sections. Consequently, quantitative morphological analysis usually requires sampling of selected sections or regions of interest and subsequent extrapolation. Extrapolation inherently introduces bias to quantitative outcome metrics. Particularly in the analysis of sparsely and non-uniformly distributed structures, such as neuromuscular junctions in skeletal muscle, relevant information in other sections may be missed, and morphometrics may lose precision and accuracy. Robust techniques that enable the detailed analysis of entire intact organs thus may improve the reliability of morphometrics and at the same time provide morphological data on an unprecedented scale.

Recent advancements in laboratory tissue processing techniques now enable researchers to render entire organs completely transparent in a chemical process termed optical tissue clearing (Richardson & Lichtman, 2015). In conjunction with the expression of fluorescent proteins in transgenic animal models or post-mortem immunostaining, target structures in cleared organs can be visualized in a preserved threedimensional tissue architecture (Belle et al., 2017; Erturk et al., 2011; Hama et al., 2011; Neckel, Mattheus, Hirt, Just, & Mack, 2016; Renier et al., 2014). The tissue clearing process first harmonizes the refractive index of the processed specimen by removing lipids, pigments and water and then matches the refractive index with an imaging solution to minimize light scattering (Erturk et al., 2011; Richardson & Lichtman, 2015). The achieved tissue transparency enables the almost unrestricted penetration of light and thus allows for fluorophore excitation even in deep tissue layers that were previously inaccessible in intact tissue (Richardson & Lichtman, 2015).

Over the last decade optical tissue clearing techniques have largely transformed neuroscience of the brain by enabling brain-wide studies of the rodent connectome (Ertürk & Bradke, 2013; Hama et al., 2011; Neckel et al., 2016; Qi et al., 2019; Ueda et al., 2020), Yet their application in experimental nerve regeneration research is presently rare. Here we describe how we adapted and applied previously developed, solvent-based optical tissue clearing techniques to enable comprehensive morphological assessments in nerve repair studies.

## Results

### Retrograde labelling studies in intact spinal cord and dorsal root ganglia

Investigators rely heavily on the quantification of regenerated motor and sensory neurons following experimental nerve repair to evaluate the extent of nerve regeneration. However, conventional tissue sectioning is labour intensive, susceptible to cutting artifacts, and requires correction factors to estimate the true number of neurons (Abercrombie, 1946). To overcome these challenges, we used a modified FDICSO protocol to clear intact spinal cord and entire dorsal root ganglia (DRGs) of adult rats and to quantify fluorescently labelled neurons via automated image segmentation.

Using immersion in a series of commercially available solvents, we achieved almost complete transparency for both, spinal cord and DRGs within 24 h (Figure 3). Thereafter, the immersed specimens were imaged via fluorescence microscopy. We found light-sheet microscopy to be best suited to large, cleared tissue specimen, due to the rapid image acquisition, minimal photobleaching and high spatial resolution (Figure 2 A to C).

**Figure 1:**
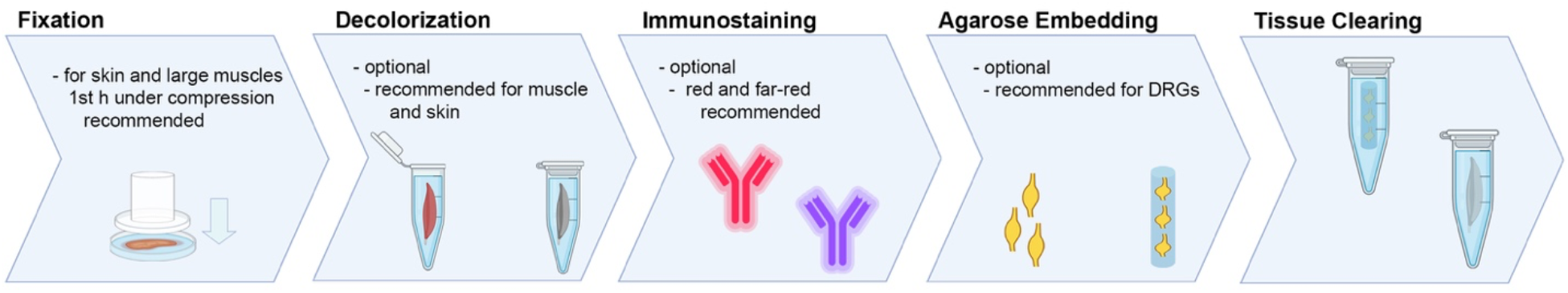
Tissue processing steps for optical clearing. After fixation by immersion in paraformaldehyde, an optional decolorization step for tissues that include high amounts of pigments (such as hemo- and myoglobin) is followed by whole organ immunostainings, preferably with red or far-red fluorophores. To streamline the image acquisition process, small organs can be embedded in agarose gel and imaged together. Thereafter, the samples undergo optical clearing in pH 9.0 adjusted and precooled (4°C) solvents in the dark. Then the cleared samples can be imaged or stored in the imaging solution, depending on the fluorophore stability. (DRG – dorsal root ganglia)

As compared to manual cell counts in tissue sections, three-dimensional imaging of the entire spinal cord and intact DRGs allows for precise quantitation of all labelled neurons. However, manual annotation of labelled cells in large tissue volumes is time-consuming. To streamline the quantitative analysis of the terabyte-sized data sets, we determined the applicability of automated image segmentation based on cell body dimensions and fluorescence intensity thresholds (Figure 2 D to F and I to L). First, we manually tagged retrogradely labelled and unlabelled (autofluorescent) neurons in the spinal cord and DRGs (L1 to L6) of three adult rats that had previously undergone retrograde labelling of the left common peroneal nerve with the fluorescent tracer Fluoro-Gold. We measured the average voxel fluorescence intensity per cell body of the Fluoro-Gold labelled neurons (n=1792 motoneurons; n=3421 sensory neurons) and randomly selected unlabelled, autofluorescent neurons (n=1120 motoneurons; n= 1774 sensory neurons, Figure 2 G and M). The fluorescence intensity of the labelled motoneurons was significantly higher compared to the autofluorescence intensity of unlabelled motoneurons (673.5 ± 109.3 vs. 385.3 ± 37.5; *p* < 0.001). Similarly, the Fluoro-Gold labelled sensory neurons in cleared DRGs had a significantly higher fluorescence intensity than the unlabelled cell bodies (1774.8.2 ± 506.5 vs. 533.3 ± 52.9; *p* < 0.001). We then used receiver operating characteristic (ROC) analyses to determine the diagnostic performance of automated segmentation of labelled neurons amongst different experimental animals based on a consistent fluorescence intensity threshold. We used the manual segmentations as a standard and identified a mean fluorescence intensity per cell of 481.4 for to be the optimal cut-off value for discriminating Fluoro-Gold labelled motoneurons from autofluorescent neurons (sensitivity: 99.3%; specificity of 98.7%; ROC area under the curve 0.9996, Figure 2 H). For sensory neurons we found even higher discriminatory accuracy using an intensity threshold of 872.1 (sensitivity: 100%; specificity of 100%; ROC area under the curve 1.0, Figure 2 P), indicating complete differentiation between the labelled neurons versus the unlabeled (autofluorescent) cell populations. The higher fluorescence intensities of DRG neurons are most likely a result of the smaller tissue volume compared to spinal cords and thus less light scattering during fluorophore excitation and emission.

**Figure 2:**
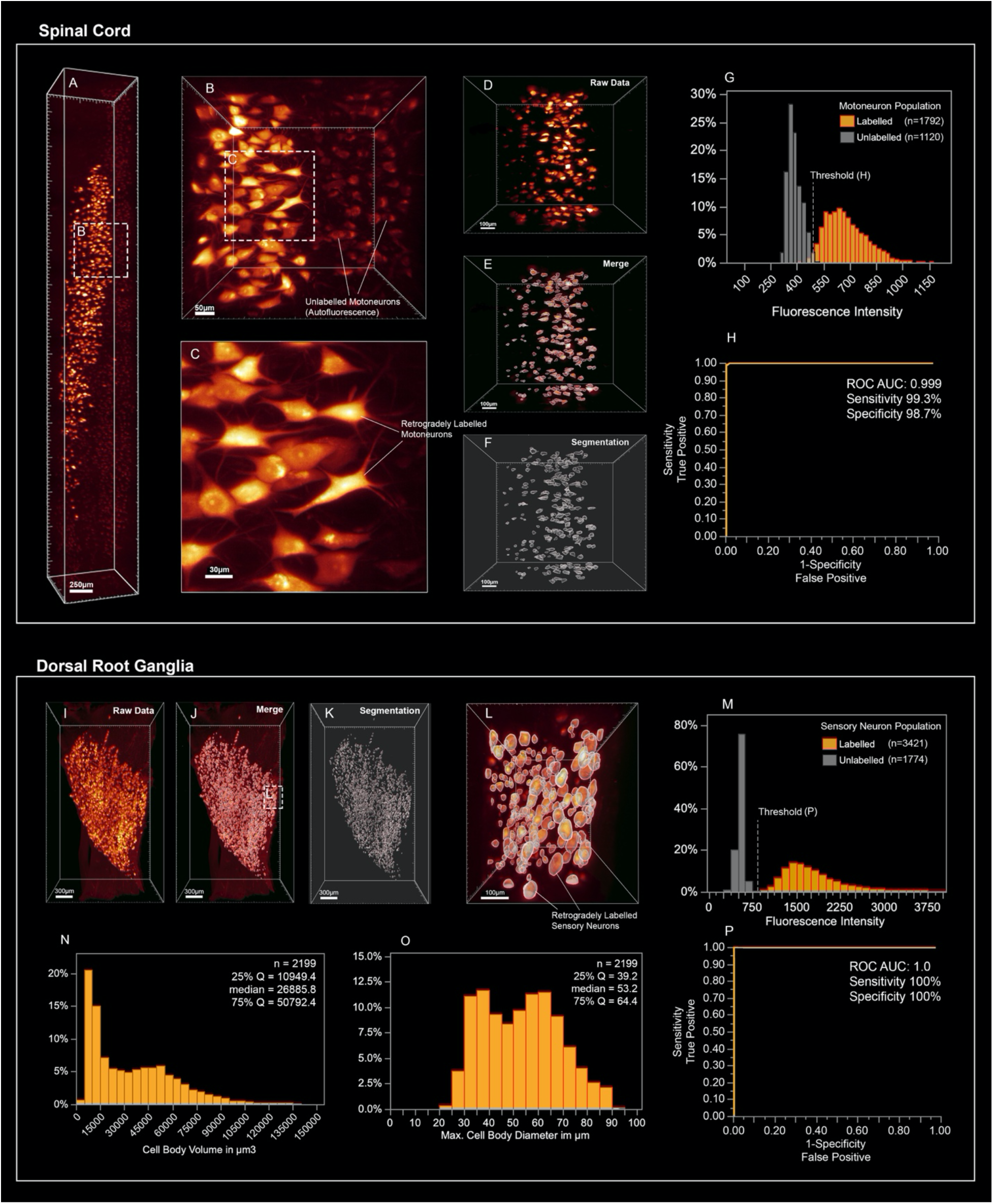
Neuron segmentations in optically cleared spinal cord and dorsal root ganglia. **(A)** Retrogradely labelled motoneurons in the ventral horn that project their axons in the common peroneal nerve. **(B and C)** Higher magnifications of labelled and autofluorescent cell bodies. **(D to F)** Automated motoneuron segmentations. **(G)** The significantly different mean voxel fluorescence intensity per cell of labelled and autofluorescent neurons enables precise, automated discrimination of labelled and unlabelled motoneurons via fluorescence intensity thresholds (ROC curve in **H**). **(I)** Optically cleared L4 dorsal root ganglion after retrograde labelling of the common peroneal nerve. A segmentation of the same DRG is shown in **(J)** and **(K)** and in higher magnification in **(L)**. **(M and P)** Similar to motoneurons, labelled sensory neurons can be identified based on fluorescence intensity with high precision and accuracy. Such automated segmentations of fluorescently labelled sensory neurons provide population-wide metrics including cell body volume in **(N)** and maximum cell body diameter **(O)** showing two histogram maxima that may indicate labelling of at least two morphologically distinct neuronal subpopulations. (ROC – receiver operating analysis; AUC – area under the curve; Q – quartile)

We then compared the neuron counts in optically cleared spinal cord and DRGs against the estimates derived from conventional cell counting technique in serial tissue sections in age matched control rats (n=3). Without correction for split cells, the number of labelled motoneurons that were counted in tissue sections significantly exceeded the neuron number in optically cleared spinal cord (697 ± 16 vs. 569 ± 23; *p* = 0.001). Similarly, the mean number of sensory neurons counted in sections of DRG L1 to L6 was substantially higher compared to the counts in cleared DRG’s (6138 ± 345 vs. 2632 ± 94; *p* < 0.001). To correct for double counted, split cells in tissue sections, we applied the Abercrombie correction factor (Abercrombie, 1946). We calculated a correction factor of 0.56 for motoneuron counts and 0.36 for sensory neuron counts based on median cell body diameters of 38.5 μm for motoneurons (n = 375 measured cell bodies) and 34.9 μm for sensory neurons (n = 400 measured cell bodies). Compared to the corrected cell counts, we identified 46% more labelled motor (*p* < 0.001) and 19% more sensory neurons (*p* = 0.009) in optically cleared, intact tissue, indicating a significant overcorrection by the Abercrombie factor (Figure 3 B and C). We further noted an approximate 50% reduction of tissue processing time in the tissue clearing group compared to the conventional technique previously used in our laboratory (Figure 3 A). Those time savings were primary attributed to the superfluous cryoprotection and labour-intensive steps such as embedding, cryosectioning and manual cell counting as part of the conventional protocol.

**Figure 3:**
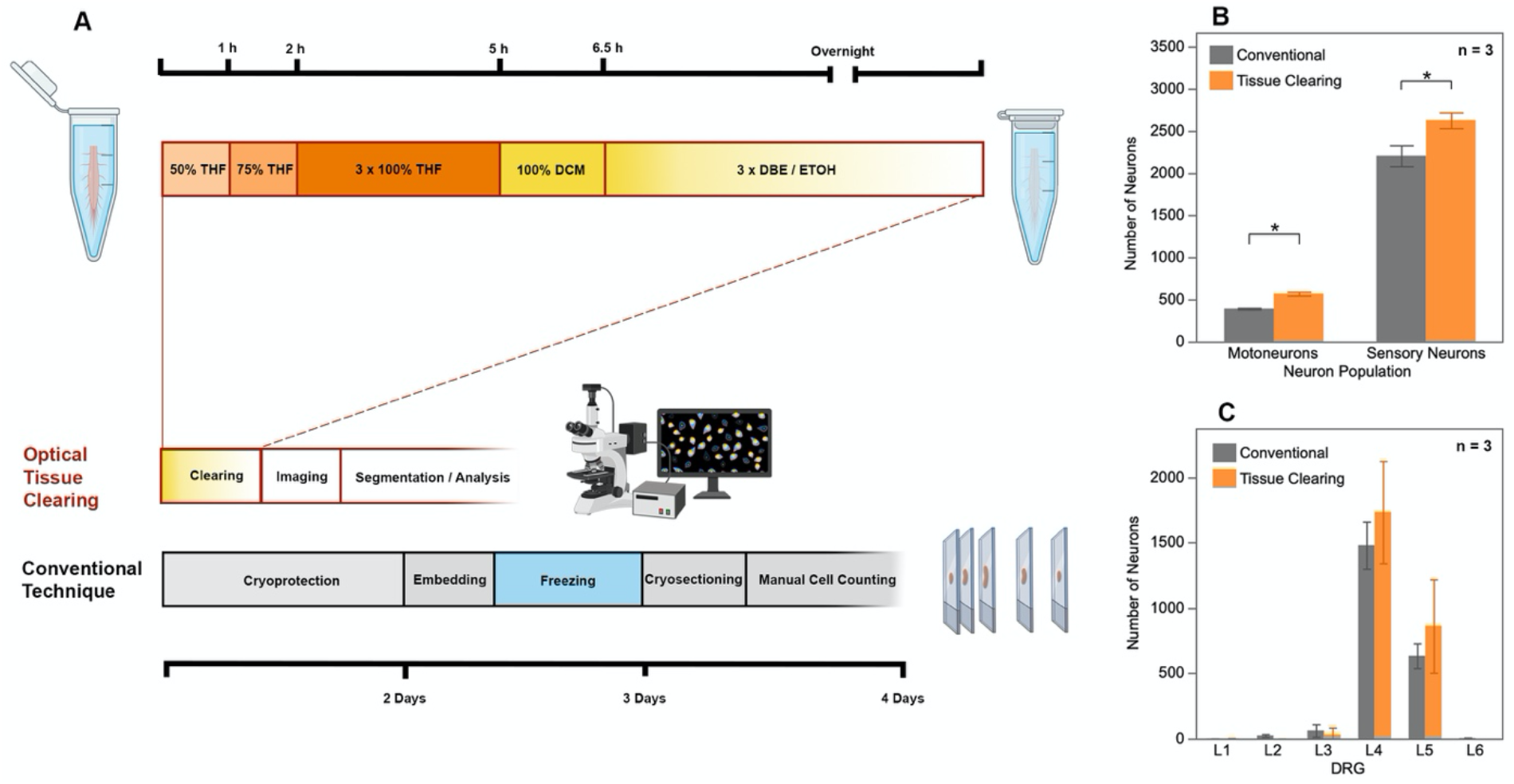
Comparing optical tissue clearing and the conventional cryosection technique for the quantitative assessment of retrogradely labelled common peroneal nerve neurons. **(A)** Processing steps for fixed tissue in both techniques, indicating substantial time savings when tissue clearing is used compared to cryosections. **(B)** Comparing the number of motor and sensory neurons derived from optically cleared (orange) and cryosectioned tissue (grey), showing that significantly more neurons were identified in intact, cleared spinal cord and DRGs. **(C)** The number of labelled common peroneal nerve sensory neurons that have been identified by both techniques in the dorsal root ganglia from L1 to L6. The results indicate a tendency of the conventional technique to overestimate cell counts in ganglia with few labelled neurons (L2 and L3) and to underestimate neuron counts in ganglia with a high density of labelled cells (L4 and L5), although the differences did not reach statistical significance in this experiment (*p* > 0.05). (THF – tetrahydrofuran; DCM – dichlormethane; DBE – dibenzyl ether; DRG – dorsal root ganglia)

Three-dimensional image segmentation of cleared organs enables a wide range of quantitative assessments of fluorescently labelled cell populations beyond automated enumeration. Possible measurements include, but are not limited to, volumetric or dimensional measurements of single cells or entire cell populations, as demonstrated in Figure 2 N and O. Here we measured the cell body volume (median 26 885.8 μm^3^) and the maximum cell diameter (median 53.2 μm) of retrogradely labelled sensory neurons that project their axons in the common peroneal nerve. As shown in the histograms of Figure 2 N and O, multiple local maxima can be distinguished indicating inclusion of morphologically distinct cellular subpopulations. Application of previously established population thresholds may allow for detailed differentiation of those subpopulations in optically cleared neural ganglia.

### Mapping the neurovascular network in optically cleared peripheral nerves

Sufficient blood supply is essential to clear debris and establish a pro-regenerative environment following peripheral nerve injury. Accordingly, ischemia has been described as a determining pathomechanism in neuropathies and a key length-limiting factor for acellular nerve allografts and synthetic conduits (Best, Mackinnon, Midha, Hunter, & Evans, 1999; Goncalves et al., 2017; Ostergaard et al., 2015). However, the historically challenging assessment of the neurovascular network in three dimensions has been a limiting factor for conclusive morphological studies. We used optical tissue clearing techniques to render peripheral nerve segments transparent and thereby make the intraneural vascular network accessible for high-resolution fluorescence microscopy (Figure 4 A). In specimens harvested from animals that did not undergo transcardial perfusion, vessels still contained red blood cells which are intensely auto fluorescent in the green emission spectrum. This can already be used as a broad approximation to map larger blood vessels, but in smaller capillaries, we found the distribution of red blood cells to be inconsistent. We therefore used intravenous injection of fluorescently conjugated CD31 antibody to label the endothelial walls of the entire vascular network. Using a far-red emission spectrum, we obtained consistent staining across upper and lower extremity peripheral nerves. Automated blood vessel tracing based on fluorescence intensity thresholds enabled mapping of the neurovascular network including capillaries with an inner diameter below that of red blood cells (< 6 μm) as shown in Figure 4 B, C and D. Beyond visualization and qualitative assessment, we were able to segment the blood vessels in inter-branch segments for detailed quantitative analysis. Exemplary, we segmented a 1000 μm segment of a rat common peroneal nerve (Figure 4). The inner diameter of the intraneural blood vessels ranged between 4.0 μm and 23.5 μm, averaging 11.6 ± 3.3 μm (Figure X D). Most inter-branch segments were short (< 40μm), but some blood vessels spanned over almost 400 μm between two branch points (Figure 4 E). When analyzing the angle between the vascular axis and the longitudinal nerve axis, the blood vessels were predominantly longitudinally oriented, as previously described (Best & Mackinnon, 1994). However, approximately 5% of the inter-branch segments were orientated perpendicular to the nerve axis, representing perforators that connect epi- and perineural to endoneurial blood vessels (Figure 4 F).

**Figure 4:**
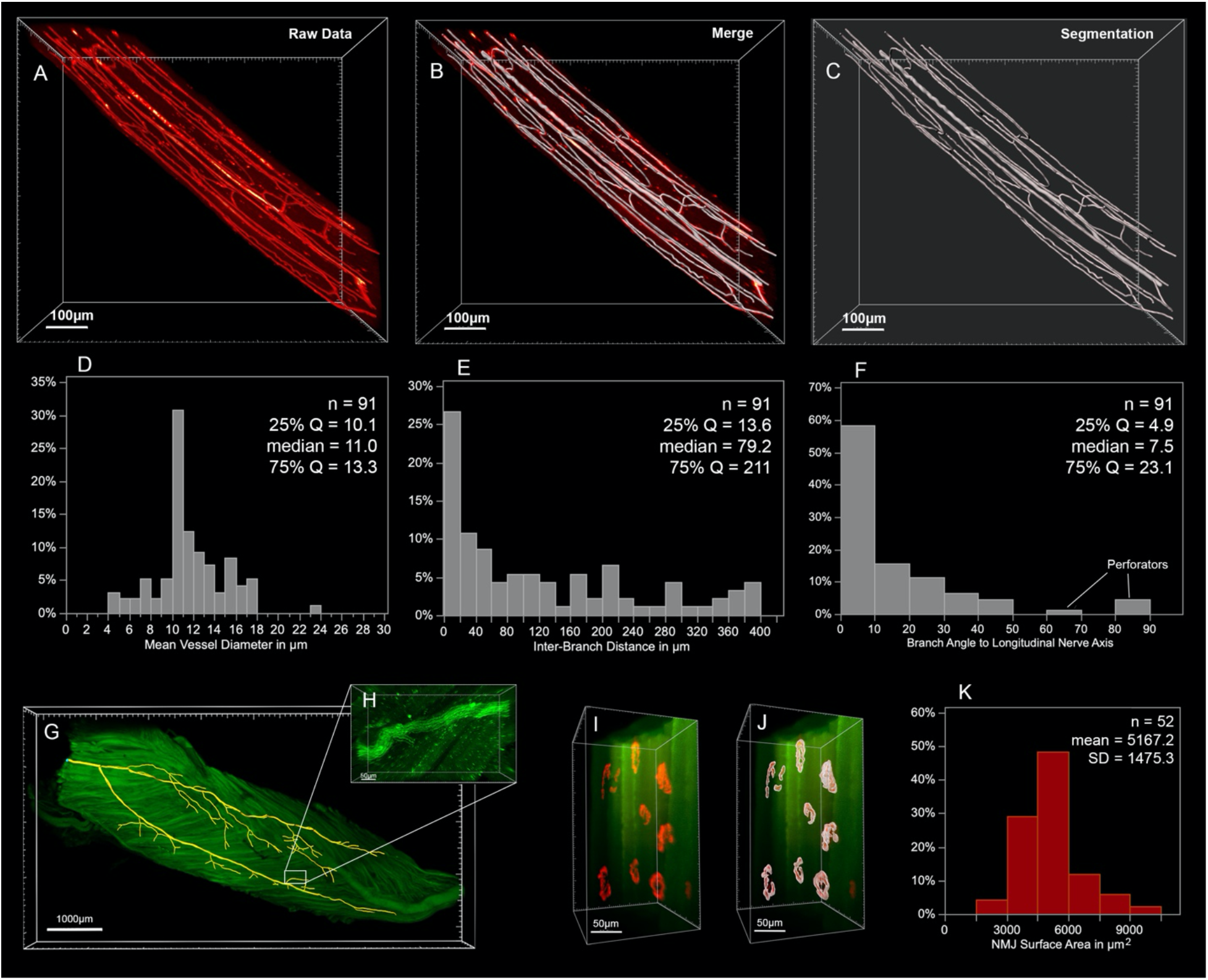
Filament tracing in optically cleared peripheral nerve and skeletal muscle. **(A)** Fluorescently labelled neurovascular system in a rat common peroneal nerve segment. Segmentations as seen in **(B)** and **(C)** can compartmentalize the blood vessel network in inter-branch segments and thereby allow for detailed analysis of the blood vessel diameter **(D**), the distance between two branching sites **(E)** or the angle of the vascular axis **(F). (G and H)** Similar fluorescence-based filament tracing techniques can be used to map the intermuscular nerve branches in the peroneus longus muscle of a transgenic thy-1 gfp+ rat with green-fluorescent axons. **(I and J)** Neuromuscular junctions stained with fluorescently conjugated α-Bungarotoxin (red) can be mapped in cleared muscle and quantitatively analyzed for their three-dimensional surface area **(K)** to determine structural changes related to development or disease. (Q – quartile; SD – standard deviation; NMJ – neuromuscular junction)

### Tracing neural projections in target organs

Transparent skeletal muscles allow for comprehensive analysis of intramuscular neuronal projections and their neuromuscular junctions. This may be used to quantitatively assess the re-innervation process following peripheral nerve repair or determine disease-related changes in the neuromuscular architecture. We used threshold-based filament tracing to visualize motor nerve branches within a peroneus longus muscle of a thy-1 gfp+ rat, a commonly used target muscle for experimental nerve repair studies (Figure 4 G and H). Notably, paraformaldehyde (PFA) fixed skeletal muscle features a pronounced autofluorescence in the green and red emission spectrum (Figure 4 H), making far-red the preferred fluorophore for muscle imaging due to a higher signal to noise ratio, as previously described (Williams et al., 2019). To visualize the neuromuscular junctions (NMJs), we intravenously injected α-Bungarotoxin conjugated with a fluorescent marker in the far red-spectrum (Chen et al., 2016). NMJ morphology is remarkably stable in rodents with intact nerves (Balice-Gordon & Lichtman, 1990). In response to denervation or reinnervation, however, NMJ’s undergo substantial structural reorganization and change their appearance and surface area (Rich & Lichtman, 1989). Such metrics can be precisely quantified on a large scale in optically cleared skeletal muscle. To demonstrate their applicability, we randomly segmented a region of interest measuring 0.5 x 0.5 x 1.0 mm within a rat peroneus longus muscle within which we identified a total of n = 52 separate neuromuscular junctions. Using the Imaris surface tool, we created surfaces with 0.3 μm surface detail for each individual NMJ to quantify the three-dimensional surface area (Figure 4 I and J). The surface area of the physiologically innervated NMJ’s ranged from 2840.3 μm^2^ to 9712 μm^2^ with an average area of 5167.2 ± 1475.3 μm^2^ (Figure 4 K).

### Dermatome-wide cutaneous innervation studies in cleared skin

To determine the applicability of optical clearing techniques to assess skin innervation, we harvested full thickness skin flaps from the sural nerve dermatome of adult Sprague Dawley rats. Immediately before harvest, we applied hair removal cream (Veet, Reckitt Benckiser, UK) to remove the strongly autofluorescent hairs. To facilitate imaging, we flattened the skin flaps with gentle pressure during the first hour of PFA fixation. Subsequently, the samples were decolorized, permeabilized and immunostained against neuronal cell markers. Using optical tissue clearing, the full thickness skin flaps were rendered transparent and then imaged via confocal microscopy. Despite moderate autofluorescence in the red spectrum, the cutaneous nerve branches were clearly visible (Figure 5 A and B). Within each nerve branch, individual axons were distinguishable that terminated in free- or lanceolate nerve endings in the skin and around hair follicles (Fig 5 E). To quantitatively assess the cutaneous innervation, we segmented a region of interest (ROI) of 2500 μm x 2000 μm (Figure 5 B to D). The skin innervation density, defined as nerve branch length per ROI area, was 9508.9 μm / mm^2^. Of note, this is 2.6-fold lower compared to the nerve fiber density in the cornea of the same rat strain (Catapano, Antonyshyn, Zhang, Gordon, & Borschel, 2018). The median branching angle of the cutaneous nerve fibers was 33.9° and the median branch length was 63.85 μm. We found 56 free nerve endings per mm^2^, some of which arose from nerve branches with more than 60 branching points within the ROI, demonstrating the arborized architecture of the cutaneous nerve plexus (Figure 5 F).

**Figure 5:**
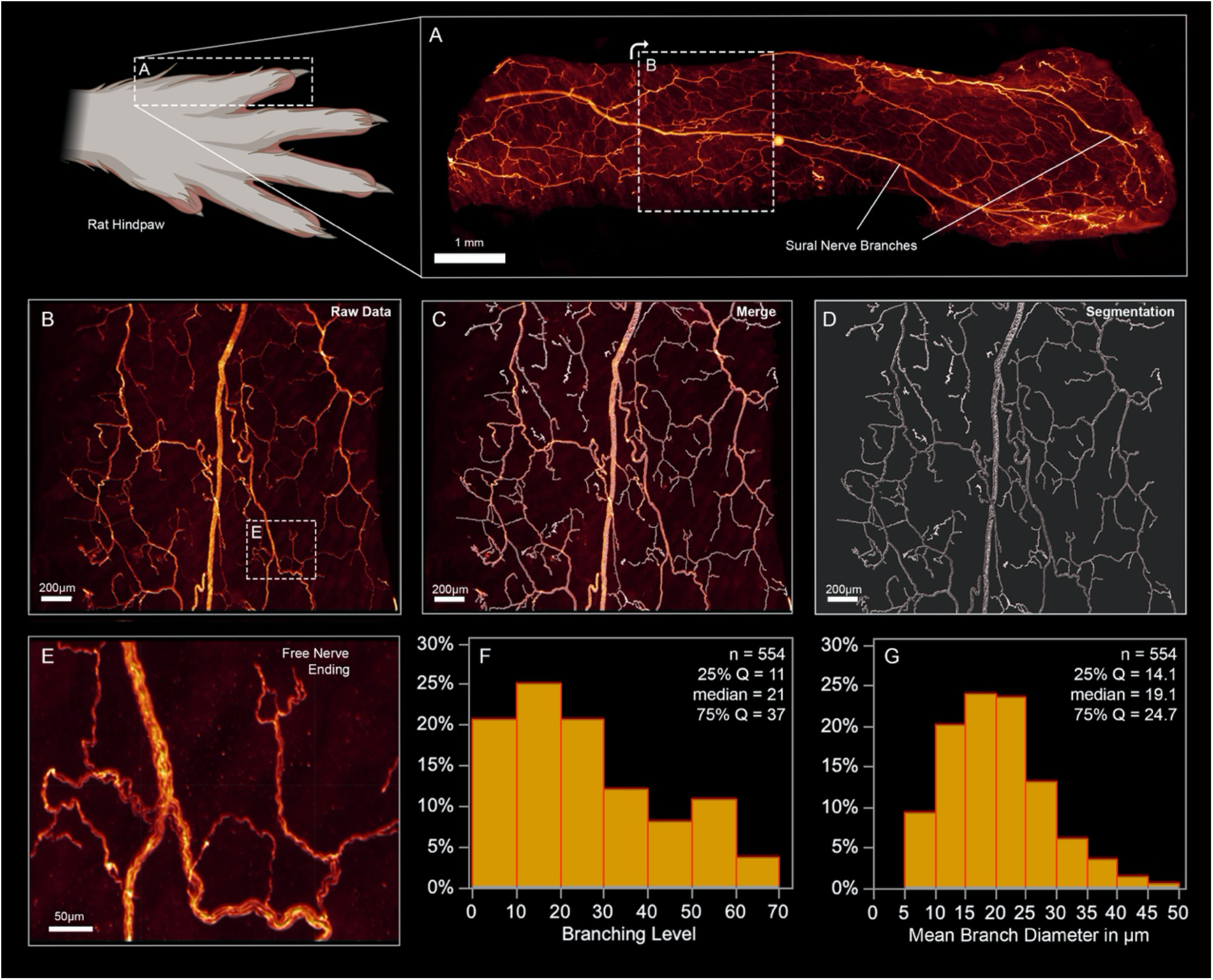
Cutaneous innervation in the rat sural nerve dermatome. **(A)** A full thickness skin graft from the dorsum of the fifth toe was harvested, immunostained against the neuronal marker beta-3-tubulin (red) and optically cleared for fluorescence imaging. **(B and E)** Cutaneous nerve branches in higher magnification. **(C and D)** Segmentations of the cutaneous innervation. The histogram in **(F)** displays the percentage frequency of the branching levels of the cutaneous nerve plexus, after compartmentalization into inter-branch segments. The branching level describes the number of branch points proximal to a segment. **(G)** The percentage frequency of the average diameter of a nerve branch. (Q – quartile)

### Refractive index matching

Specialized microscopes are available with optics optimized for the high refractive index (1.56) of cleared tissue immersed in pure dibenzyl ether. However, widely used glycerol or CLARITY objectives are optimized a refractive index of around 1.45. The refractive index mismatch between sample and objective creates blurred images due to spherical aberration (Richardson & Lichtman, 2015). To overcome the refractive index mismatch for those objectives, we used immersion in ethanol / dibenzyl ether solutions as previously described (Carro, Paroutis, Woolside, & Harrison, 2015). Lowering the refractive index of the immersion solution, however, inevitably affects sample transparency. To optimize the imaging solution for glycerol / CLARITY objectives, we determined the best trade-off between transparency-dependent imaging depth, and spherical aberration due to refractive index mismatch in ascending ethanol / dibenzyl ether solutions. The best images were achieved with a 34.1 vol% ethanol in dibenzyl ether solution for spinal cords, and a 30.1 vol% solution for agarose embedded DRGs.

## Discussion

Here we demonstrate the applicability of optical tissue clearing to experimental nerve regeneration studies. Optical tissue clearing methods have been introduced to overcome imaging depth limitations in harvested tissue by reducing light scattering to a level of almost complete transparency (Erturk et al., 2011; Richardson & Lichtman, 2015). In conjunction with light-sheet fluorescence microscopy and powerful data processing tools, these techniques enable organ-wide morphological readouts and objective analysis with subcellular resolution. This has revolutionized the field of brain connectomics (Schneider-Mizell et al., 2016) and holds similar promise for morphological studies of peripheral nervous system in development, health and disease.

A variety of tissue clearing techniques have been developed and continuously refined, each having their respective advantages and drawbacks for different research applications (Ertürk & Bradke, 2013; Jing et al., 2018; Neckel et al., 2016; Qi et al., 2019; Renier et al., 2014). In our laboratory, we adopted a slightly modified FDISCO protocol due to its powerful clearing capabilities, rapid processing times, low-costs and easy applicability requiring only conventional laboratory equipment. We were able to process entire organs with this solvent-based technique and thereby overcame the risk of cutting artifacts in precious experimental samples. The protocols have been proven to reliably provide high levels of transparency for all tested organs. When the level of transparency was insufficient due to application errors or following long-term storage, immersion in 100% THF can simply be repeated even months after the initial clearing process to readjust the refractive index and achieve excellent transparency.

Adopting optical tissue clearing techniques in our laboratory has increased time efficiency by overcoming labour-intensive steps such as cryosectioning and manual cell counting. This methods also avoids timeconsuming transcardial perfusion with PFA and instead uses hydraulic extrusion of the spinal cord (Richner, Jager, Siupka, & Vaegter, 2017). Beyond substantially increased time-efficiency, this method does not require dissections of PFA-soaked animals and thus drastically decreases the associated risk of accidentally damaging the spinal cord and avoids exposure to hazardous fumes.

For retrograde labelling studies, the inconsistent, multisegmental distribution of sensory neurons that project their axons into a peripheral nerve necessitates examination of multiple dorsal root ganglia (DRGs) per experimental animal to reliably capture all neurons (Figure 3 C). When using conventional tissue sectioning techniques, each DRG represents hours of work. Investigators consequently limit the number of examined specimen to one or two ganglia per animal (e.g. L4 and L5 for common peroneal nerve studies) (Tajdaran, Chan, Shoichet, Gordon, & Borschel, 2019; Zuo et al., 2020). Due to their time- and work efficiency (Figure 3 A), tissue clearing techniques overcome these limitations. To further streamline the imagine acquisition process, we embedded all relevant DRGs of the same animal together in 1% Agarose gel columns before tissue clearing. Thereby we were able to reduce the number of required microscope setups and further increase time-efficiency.

Three-dimensional imaging with high spatial resolution in all planes allows for reliable application of automated segmentations and subsequent quantitative analysis based on objective parameters. This is in stark contrast to manual cell counting in serial tissue sections where, among cells with high fluorescence intensity, a subset of weakly stained cells often present investigators with challenges whether to count the cell or not. Such numerical insecurities are further amplified by double counted split cells and the consequent need to apply correction factors to estimate the true number of cells (Abercrombie, 1946). We observed that neuron counts derived from conventional tissue sections exceed the true number of retrogradely labelled neurons but the Abercrombie correction factor tends to overcorrect neuron counts (Figure 3 B). We have shown that automated cell counting, based on a predefined intensity threshold provides highly accurate cell counts, with a sensitivity above 99.3% and specificity greater 98.7% (Figure 2 H and P). The ROC analyses further indicated excellent performance of a consistent threshold for discriminating Fluoro-Gold labelled cells from autofluorescent cells across different experimental animals. This substantially reduces observer-introduced bias for retrograde neuronal labelling experiments and was possible by standardizing the labelling (dye concentration and exposure time), tissue processing (fixation and immersion times) and imaging procedures (laser intensity, resolution, z-step-size).

Comprehensive analysis of the re-organization of neural projections following nerve surgery, particularly nerve transfers, is critical to better understand their functional impact in rodent models and translate experimental results in well-founded clinical hypotheses. An advantage of optically cleared tissue is the preserved three-dimensional tissue architecture. Tracing filamentous structures with a high length-to-diameter ratio, such as nerve fibers, that span across in large tissue volumes has been historically challenging in tissue sections (Helmstaedter, Briggman, & Denk, 2011). The high tissue transparency achieved by optical clearing now provides the opportunity to map the entire neuromuscular connectome of a regenerating nerve and thereby unveils an unprecedented level of information for morphological outcome assessments in experimental nerve repair studies. Particularly for the study of complex arborized networks, such as axons or blood vessels, three-dimensionality inherently provides a better understanding of spatial relations and structural coherences. Of note, we have demonstrated the capabilities of software-assisted segmentation and quantitative assessment for the neurovascular network in a rat common peroneal nerve (Figure 4). By compartmentalizing the vascular network in inter-branch segments, we were able to differentiate distinct vessel types by diameter and projection angle to quantitatively analyze the intraneural blood vessel architecture in great detail. Such morphometrics may be valuable to study the reorganization of the vascular network in long allografts or assess neurovascular pathologies.

Optical tissue clearing, in conjunction with light-sheet fluorescence microscopy, enables rapid data acquisition for large, intact tissue volumes. This allows for large-scale morphometric analysis without sacrificing subcellular resolution. For nerve repair studies, these capabilities can be leveraged for neuromuscular junction morphometry in a skeletal muscle or dermatome-wide cutaneous innervation studies. Although conventional two-dimensional analysis of arborized neural projections, such as assessing intra-epidermal nerve fiber density in skin biopsies, has been proven to be valuable in determining disease progression of systemic neuropathies, such approaches do not accurately reflect the complex neural morphology. This holds particularly true when nerve fibers are unequally distributed across the region of interest, such as during the process of skin reinnervation following nerve repair. We demonstrated that clearing techniques may be used to render large, full-thickness skin samples transparent, enabling rapid, objective and detailed morphometric readouts of entire dermatomes and thus provide unprecedented insights into the cellular activities governing cutaneous (re-)innervation. Consequently, readily applicable clearing protocols may inspire attention to the reorganization of the sensory system in future nerve repair studies, in both skin as well as any other tissue type.

Imaging intact organs, however, comes with the inherent limitation of a restricted number of target structures that can be visualized in the same sample. This is due to the limited number of distinguishable fluorescent emission spectra during image acquisition. To overcome this limitation, other, hydrogel-based techniques have been developed that allow for multiple rounds of de- and re-stainings of the same specimen (Murray et al., 2015). Another limitation that must be considered is the dehydration-associated effect of tissue shrinkage that occurs in solvent-based tissue clearing protocols. This must be considered when interpreting morphometry. Conversely, tissue shrinkage facilitates the deep imaging of large samples that otherwise would exceed the maximum working distance of high-numerical aperture objectives.

Since 2011, the field of optical tissue clearing and whole-organ immunostaining is rapidly evolving (Erturk et al., 2011). Novel machine learning models are already on the horizon and are going to streamline the further segmentation process of large, multiple terabyte-sized image stacks by reducing manual processing steps and potential observer bias. Presently developed panoptic imaging methods and compatible microscopes are soon going to enable rapid imaging of entire, optically cleared rodents (Cai et al., 2019). In conjunction with capable machine learning models and transgenic animal models (e.g. brainbow mice), this will allow for automated tracing of the entire neural connectome of the central and peripheral nervous system in the same experimental animal, likely offering exciting insights in biological processes and a deeper understanding of systemic reorganization after nerve injury and repair.

## Conclusion

We demonstrated the applicability of optical tissue clearing techniques and computational image segmentation for a variety of tissues relevant to experimental nerve repair studies and outlined their potential benefits and current limitations. Integrating those techniques into our research has increased time-efficiency while enabling objective and accurate morphological studies on an unprecedented scale. Given the advantages of optical tissue clearing over conventional methods, this technique is likely to be a key technology for future experimental studies of nerve injury and repair.

## Methods

This study is reported in accordance with the ARRIVE 2.0 guidelines (Animal Research: Reporting of *In Vivo* Experiments)(Percie du Sert et al., 2020).

### Experimental animals

We included a total of n=6 adult (250 – 300 g), female transgenic rats with a Sprague Dawley genetic background. This in-house bred transgenic strain endogenously expresses green fluorescent protein (GFP) in motor and sensory neurons under the thy-1 promotor (*thy-1* gfp+) (Magill, Moore, Borschel, & Mackinnon, 2010). All animals were housed in a central animal care facility with fresh water and pellet food *ad libitum*. A constant room temperature (22°C) and a circadian rhythm of 12h per 24h illumination was automatically maintained. All procedures were performed in strict accordance with the National Institutes of Health guidelines, the Canadian Council on Animal Care (CCAC) and were approved by the Hospital for Sick Children’s Laboratory Animal Services Committee.

### Experimental procedures

#### 1. Retrograde neuronal labelling *in-vivo*

All surgical procedures were performed under aseptic conditions and inhalational anesthesia with an Isoflurane (Baxter, Illinois, USA) oxygen mixture (3%, flow: 2 L/min). For analgesia, 4 mg/kg body weight extended-release Meloxicam (Metacam, Boehringer Ingelheim, Ingelheim, Germany) was injected subcutaneously. The rat was placed on a heat pad to maintain body temperature and the left hind leg shaved and surgically cleaned with an alcohol / betadine rub. The common peroneal nerve was exposed within anatomical planes through a dorsolateral-gluteal muscle-splitting incision and cut 5 mm distal to the sciatic bifurcation. The wound bed including the adjacent peripheral nerves were carefully covered with a sterile, fluid-repellent drape to prevent leakage of the fluorescent tracer. The end of the proximal common peroneal nerve segment was placed in a well containing 5 μl 4% Fluoro-Gold (Fluorochrome LLC, Denver, USA) in double distilled water. Sterile petroleum jelly was used to keep the nerve in place and prevent desiccation. After 60 min the wounds were thoroughly irrigated with saline, dried, and closed in three layers with 5-0 Vicryl (Ethicon, Ohio, USA) sutures. Experimental animals were recovered in a warm environment prior to returning to the housing facility.

#### 2. Tissue harvest and fixation

Seven days after the retrograde labelling procedure, the rats were euthanized and the entire spinal cord and the dorsal root ganglia (DRG) from L1 to L6 were harvested bilaterally using a hydraulic extrusion technique as previously described (Richner et al., 2017). Transcardial perfusion was not required. We further harvested the contralateral common peroneal nerve and peroneus longus muscle and a full thickness skin flap of the contralateral dorsal hind-paw. The specimens were fixed by immersion in precooled 4% paraformaldehyde (PFA) at 4°C for 24 h, then washed and stored in precooled PBS in the dark at 4°C until further processing. Detailed laboratory protocols are included in the supplemental material. For the skin flaps, we applied light compression in a petri dish for the 1^st^ hour of fixation to flatten the sample. Similarly, flattening by compression can be used for large skeletal muscles when the imaging objective has a limited working distance.

#### 3. Immunostaining

For heavily colored specimens, such as muscle or skin, we used a decolorization step in 25% Quadrol (122262, Sigma-Aldrich, Missouri, USA) in double distilled water at 37°C for two days with daily changes of the solution prior to the immunostainings. The following antibodies were used: rabbit anti-beta 3 tubulin (18207, Abcam, Cambridge, UK; 1:200 dilution) with goat anti-rabbit Alexa Fluor 555 conjugated secondary antibody (A-21428, Invitrogen, California, USA; 1:300). For tail vein injections 1 h prior to euthanasia we used 20 μg α-Bungarotoxin conjugated with Alexa Fluor 647 (B35450, Thermo Fisher Scientific, Massachusetts, USA) or 30μg Alexa Fluor 647 conjugated CD31 Antibody (102516; Biolegend, California, USA) in 200 μl sterile saline respectively (Chen et al., 2016). We used whole-organ immunostainings as previously described (Qi et al., 2019). Briefly, after fixation and optional decolorization the samples were pretreated with in ascending methanol series (50%, 80%, 2 x 100% Methanol in double distilled H_2_O for 30 min each) followed by an overnight step in 5% H_2_O_2_ / 20% DMSO (276855, Sigma Aldrich) / Methanol at 4°C to reduce background fluorescence and increase permeability. Subsequently, the samples were treated in 100% methanol for 45 min twice followed by two 45 min steps in 20% DMSO / Methanol and a descending Methanol series (80%, 50% Methanol in double distilled H_2_O for 30 min each). Then the samples were washed in PBS for 1 h twice followed by an overnight step. Next samples were permeabilized in PBS / 0.2% Triton X-100 (X100, Sigma Aldrich) / 20% DMSO / 0.3M glycine overnight at 37°C and blocked in PBS / 0.2% Triton X-100 / 10% DMSO / 5% normal donkey serum for 3 days. Following two 2h washing steps in PTwH (PBS / 0.2% Tween-20 (P9416, Sigma Aldrich) / 0.01mg/ml Heparin) the sample was immersed in primary antibody dilutions in PTwH / 5% DMSO / 3% normal donkey serum at 37°C for 2-4 days, followed by a 2-day wash in PTwH on a shaker with four changes of the washing solution. Then the samples were incubated in secondary antibody diluted in PTwH / 3% normal donkey serum for 2-4 days at 37°C followed by a 2-day wash in PTwH on a shaker with at least four changes of the washing solution.

#### 4. Optical tissue clearing

For optical tissue clearing we used a modified FDISCO protocol (Qi et al., 2019). Prior to optical clearing we embedded the DRGs in PBS / 1% Agarose gel columns in one plane to facilitate subsequent image acquisition. For optical clearing, the tissue was immersed in an ascending series of precooled (4°C) Tetrahydrofuran (186562, Sigma-Aldrich) double distilled water solutions (50 vol%, 75 vol%, 3 x 100 vol%) for dehydration and delipidation. Triethylamine (471283, Sigma-Aldrich) was used to adjust the pH of the solutions to 9.0 and all steps were performed in the dark at 4°C on an orbital shaker (Qi et al., 2019). For further delipidation of particularly lipid-rich specimen such as spinal cord, we used a subsequent step in 100 % Dichloromethane (270997, Sigma-Aldrich) at room temperature. The immersion time ranged between 30 min and 2h for each step, depending on the tissue type and specimen size (See supplemental material). Then the specimen was immersed in Ethanol / Dibenzyl ether (108014, Sigma-Aldrich) solutions overnight at 4°C in the dark, with two changes of the solution, to match the refractive index of the tissue with the microscope objective. To optimize the imaging solution for glycerol / CLARITY objectives, we determined the best trade-off between transparency-dependent imaging depth and spherical aberration in ascending ethanol / dibenzyl ether solutions for each tissue.

#### 5. Imaging, data processing and analysis

For image acquisition of cleared spinal cord and DRGs we used a 3D light-sheet fluorescence microscope (Zeiss Lightsheet Z.1, Carl Zeiss Microscopy GmbH, Jena, Germany) equipped with a p.co edge 5.5 Camera (PCO AG, Kehlheim, Germany) and a 20x / 1.0 CLARITY objective optimized for a refractive index of 1.45. For Fluoro-Gold labelled specimen we used a 405 nm (20mW) laser and Zeiss Zen Lightsheet 2014 software (Carl Zeiss Microscopy GmbH, Jena, Germany). To enable automated batch processing and comparative analysis, we used consistent illumination plane settings and laser intensities of 2.5%.

Damage to the objective was prevented by exposure to dibenzyl ether, we used borosilicate glass tubing (Kavalierglass A.S., Křížová, Czech Republic) to separate the immersed specimen from the microscope optics. This tubing matches the refractive index of the adjusted tissue (1.472) and is resistant to the dibenzyl ether (Carro et al., 2015). For cleared peripheral nerves, skeletal muscle and skin flaps we used a Leica SP8 Lightning confocal microscope (DMI8, Leica Microsystems, Wetzlar, Germany), equipped with Leica LAS software, a Hybrid detector (HyD) and a p.co Edge 5.5 camera (PCO AG, Kehlheim, Germany). We used a 20 x / 0.75 (W) and a 40 x / 1.3 (W) objective with the following lasers: 405 nm (50 mW), 488 nm (20 mW), 552 nm (20 mW), 638 nm (30 mW). For inverted confocal microscopy, we used Cell Imaging Dishes with a 1.0 μm thin cover glass bottom (Eppendorf AG, Hamburg, Germany) to leverage the full working distance of the objective for maximum imaging depth.

For data processing, we used arivis Vision 4D (version 3.0, arivis AG, Rostock, Germany) or Leica LAS (Leica Microsystems, Wetzlar, Germany) for three-dimensional tile stitching, Huygens deconvolution software (Scientific Volume Imaging, Hilversum, Netherlands) and Imaris (Version 9.5.1, Bitplane AG, Zurich, Switzerland) for image segmentation, fluorescence measurements and quantitative morphometric analyses. Selected images in this publication are created with BioRender.com.

#### 6. Conventional tissue processing and manual quantification of retrogradely-labelled neurons

To validate the software-based neuron counting in optically cleared spinal cord and DRGs we compared the results against the conventional enumeration technique in serial cryosections. Therefore, we performed Fluoro-Gold retrograde labelling on three randomly selected rats and harvested the tissue with the abovementioned techniques. The specimens were fixed in 4% PFA at 4°C for 48 hours and then in 4% PFA with 30% sucrose for cryoprotection for 2 - 5 days at 4°C. The Spinal cord and DRGs (L1 to L6) were then embedded in Tissue Freezing Medium (Electron Microscopy Sciences, Hatfield, USA) and frozen at – 80°C in for a minimum of 12 hours. Then the tissue blocks were mounted in a cryostat microtome (CM3050S, Leica Microsystems, Wetzlar, Germany) and cut in 50 μm longitudinal sections (spinal cord) or 20 μm transverse sections (DRG) as previously described (Zuo et al., 2020). The tissue sections were mounted on Superfrost Plus microscope glass slides (Fisher Scientific, Pittsburgh, PA, USA) and examined under an epifluorescence microscope (Leica DM2500, Leica Micosystems, Wetzlar, Germany) at 10x magnification. A blinded examiner counted the Fluoro-Gold-labelled neurons in every spinal cord section and every fifth DRG section (Zuo et al., 2020). The total motoneuron counts were calculated with the widely used Abercrombie correction factor to account for double counted, split neuronal cell bodies that partially lie in two adjacent sections (Abercrombie, 1946).

#### 7. Statistical analysis

For statistical analysis we used JMP (version 15.1.0, SAS Institute, Cary, USA). Descriptive statistics were calculated, and means are expressed with standard deviations. To test for normality of continuous variables, we used normal quantile plots and Anderson-Darling tests. For pairwise comparison of normally distributed, continuous data we used two-sided Student t tests. For non-normally distributed variables we used Wilcoxon tests. To determine the predictive ability of fluorescence intensity thresholds to discriminate neuron populations, we used receiver operating characteristic (ROC) analyses. A significance level of 5% was used (*p* < 0.05)

